# Linking FANTOM5 CAGE peaks to annotations with CAGEscan

**DOI:** 10.1101/126474

**Authors:** Nicolas Bertin, Mickaël Mendez, Akira Hasegawa, Marina Lizio, Imad Abugessaisa, Jessica Severin, Mizuho Sakai-Ohno, Timo Lassmann, Takeya Kasusawa, Hideya Kawaji, Yoshihide Hayashizaki, Alistair R. R. Forrest, Piero Carninci, Charles Plessy

**Affiliations:** RIKEN Center for Life Science Technologies, Division of Genomics Technologies, Japan; RIKEN Omics Science Center, Japan; Present address: Human Longevity Singapore Pte. Ltd., Singapore; Present address: Department of Computer Science, University of Toronto, Canada; Present address: Telethon Kids Institute, The University of Western Australia, Australia; RIKEN Preventive Medicine and Diagnosis Innovation Program, Japan; Present address: Harry Perkins Institute of Medical Research, QEII Medical Centre and Centre for Medical Research, the University of Western Australia, Australia

## Abstract

The FANTOM5 expression atlas is a quantitative measurement of the activity of nearly 200,000 promoter regions across nearly 2,000 different human primary cells, tissue types and cell lines. Generation of this atlas was made possible by the use of CAGE, an experimental approach to localise transcription start sites at single-nucleotide resolution by sequencing the 5′ ends of capped RNAs after their conversion to cDNAs. While 50% of CAGE-defined promoter regions could be confidently associated to adjacent transcriptional units, nearly 100,000 promoter regions remained gene-orphan. To address this, we used the CAGEscan method, in which random-primed 5′-cDNAs are paired-end sequenced. Pairs starting in the same region are assembled in transcript models called CAGEscan clusters. Here, we present the production and quality control of CAGEscan libraries from 56 FANTOM5 RNA sources, which enhances the FANTOM5 expression atlas by providing experimental evidence associating core promoter regions with their cognate transcripts.

## Background & Summary

CAGE (Cap Analysis Gene Expression, [1]) is the method of choice for studying gene regulation through quantitative analysis of transcription start sites (TSS, sequence ontology term 0000315) [2]. By sequencing the 5′ end of cDNA-converted capped RNAs, CAGE enables the identification of core promoter regions and 5′ end transcriptional activity. Large scale application of CAGE by the FANTOM consortium to nearly 2,000 human RNA sources including primary cells, whole-tissue extracts and cell lines [3, 4] identified nearly 200,000 core promoter regions active within the human genome [5].

Although CAGE enables the location of TSS at a single nucleotide resolution, the determination of their connection to downstream known gene structures or to independent novel RNAs is limited to positional computational inference and low-throughput gene-by-gene experimental validations. Half (101,893/201,802) of the FANTOM5’s active core promoter regions did not co-localize within a reasonable distance with 5′ termini of annotated gene models. To experimentally associate these orphan core promoter regions to transcriptional units, we employed *CAGEscan* [6], an approach in which paired-end sequencing of the 5′ end of cDNA-converted capped RNAs with their cognate randomly priming sites enables the unequivocal association of individual TSS to transcripts exons. In a previous project, focused on analysing the translatome of Purkinje neurons in rat [7], the CAGEscan approach annotated 43 % of the core promoters active in rat’s Purkinje neurons that we detected but had no by direct overlap with Ensembl transcripts.

Here, we selected 56 RNA sources which upon FANTOM5 CAGE profiling revealed the greatest levels of transcriptome diversity and prepared individual CAGEscan libraries, with 6 of these 56 RNA sources prepared in duplicate (see Table 1). Using the FANTOM5 core promoter atlas as seed, we clustered the CAGEscan paired-end reads in a collection of 112,315 models called *CAGEscan clusters*. To de-orphanise FANTOM5 promoters, we intersected the CAGEscan clusters with GENCODE 18 gene models. Of the 85 % that intersected, 33,632 clusters had no annotation in FANTOM5, thus revealing novel and alternative promoters to known genes. We made these data available along with the FANTOM5 CAGE atlas data, as well as ready for manual inspection and analysis via the ZENBU genome browser [8] (see Figure 1 and Data Citation 1).

**Table 1:**
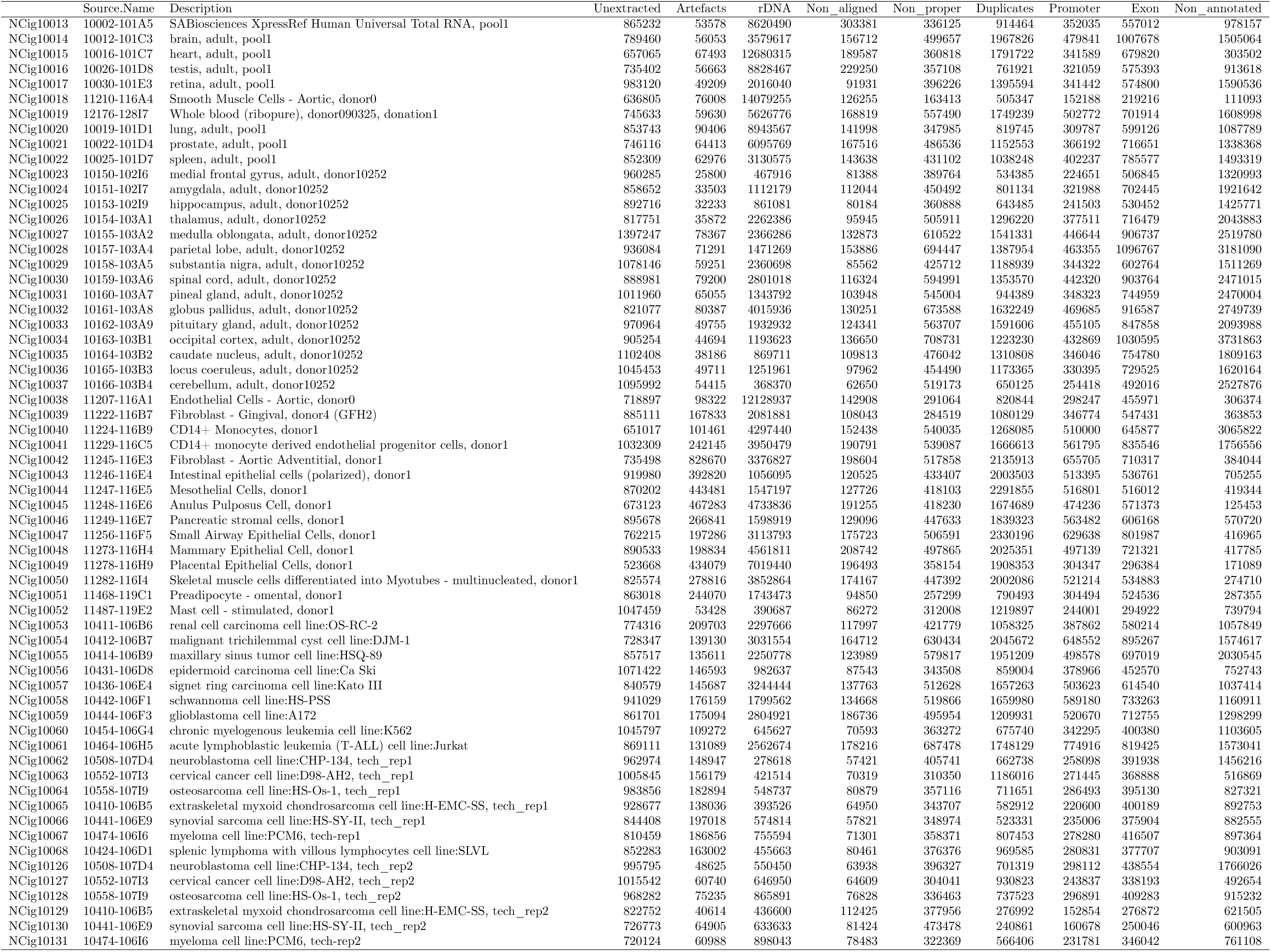
Summary of the libraries prepared. The RNA identifier (Source.Name) can be searched in the FANTOM5 SSTAR database [9, 10]. The RNA samples are also described in the SDRF files distributed alongside the FASTQ sequences and alignments, as well as the raw alignment statistics.

**Figure 1:**
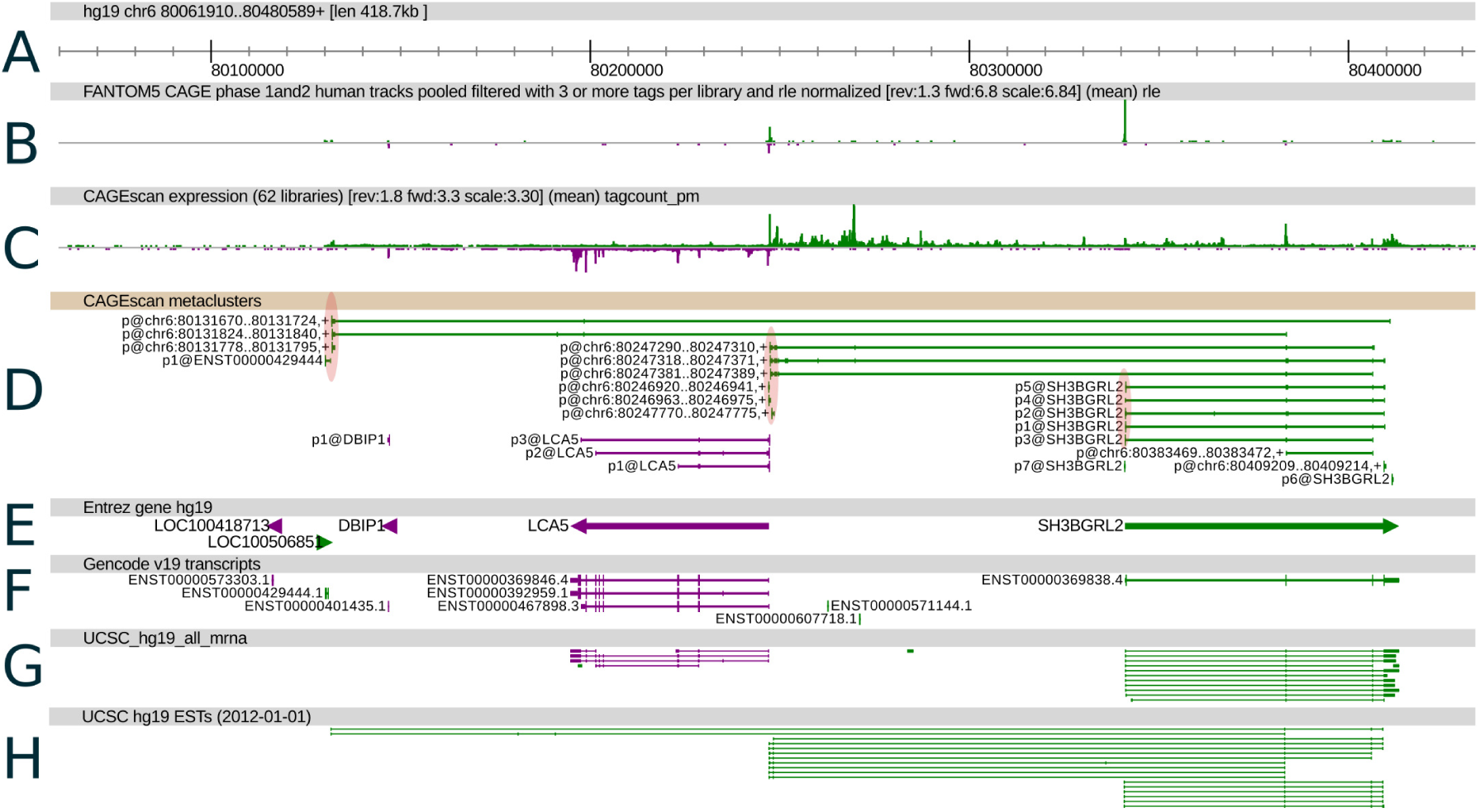
CAGEscan clusters revealing new promoters for the SH3BGRL2 gene. Features on the plus and minus strand are displayed in green and purple respectively. Promoter regions of interest are highlighted with ellipses in track D. A: Genomic coordinates. B: FANTOM5 CAGE signal as a quantitative histogram. C: CAGEscan CAGE signal. D: CAGEscan meta-clusters, combining pairs for all libraries. The name of the seed CAGE peak is indicated on the left of each cluster. E: NCBI Gene bodies. F: GENCODE 19 annotations. G: GenBank mRNA sequences. H: EST sequences supporting the CAGEscan clusters.

## Methods

All human samples used in the project were either exempted material (available in public collections or commercially available), or provided under informed consent. All non-exempt material is covered under RIKEN Yokohama Ethics applications (H17-34 and H21-14). The CAGEscan libraries were prepared as described earlier [11]. In brief, 500 ng of RNA were reverse-transcribed in presence of random primers and template-switching oligonucleotides, amplified by PCR and sequenced paired-end (2 × 36 nt) on Illumina GAIIx sequencers, one sample per lane. The barcode sequence GCTATA, present in every sample, acted as the spacer that we introduced in [12] to decrease the amount of strand-invasion artifacts. The paired-end sequences were then processed with the MOIRAI workflow system [13], with a template implementing the workflow OP-WORKFLOW-CAGEscan-FANTOM5-v1.0, described below and in Figure 2.

For each pair, the first (CAGE) and second (CAGEscan) reads in FASTQ format were demultiplexed. The first 9 bases of the CAGE reads were trimmed as they contain the sample barcode and the template-switching linker. CAGEscan paired-end reads that did not contain the exact barcode and linker sequences were discarded. The first 6 bases of the CAGEscan reads were trimmed, because they originate from the random primers and not the cDNAs, and therefore are prone to errors caused by mismatches during the hybridization to the RNAs, that are well tolerated by the reverse-transcriptase [14].

**Figure 2:**
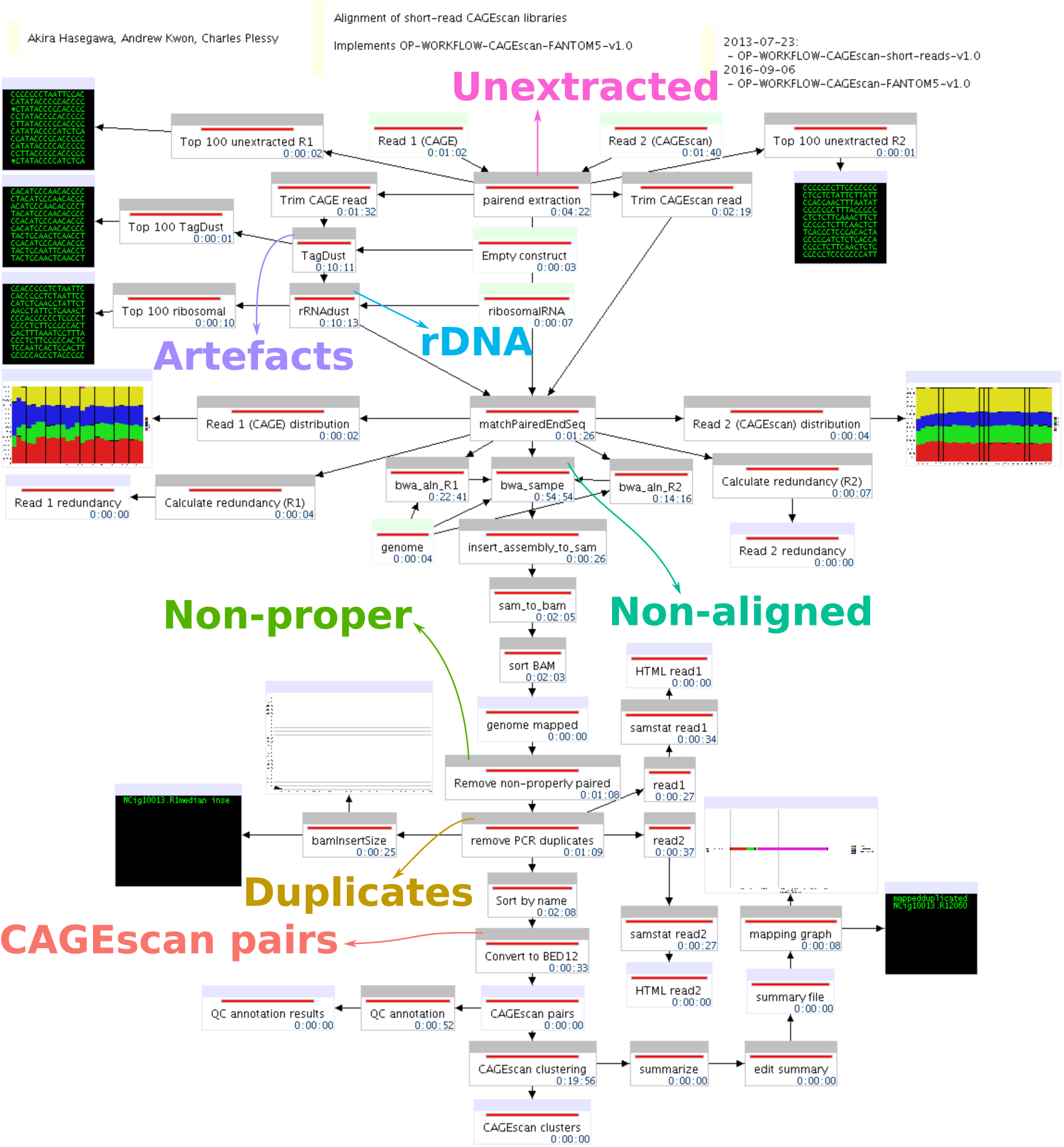
Processing pipeline. The diagram made of boxes connected by black arrows displays the MOIRAI workflow completed for one (NCig10013) of the 62 CAGEscan libraries. The colored text and arrows overlayed on the diagram represents the points where the main alignment statistics are calculated to summarize the number of read pairs passing all the filters (CAGEscan pairs) or discarded at each step of the processing pipeline (Unextracted, rDNA, Artifacts, Non-aligned, Non-proper, Duplicates).

The CAGE and CAGEscan reads were then filtered independently with the TagDust program version 1.13 [15], using the sequences of empty constructs and primers as artifact library. They were then compared to reference sequences of ribosomal genes (GenBank: U13369.1) using the rRNAdust program version 1.03. Reads whose mates were discarded by these two filters were then removed.

FASTQ formatted cleaned paired-end reads were then aligned on the human genome version hg19 with BWA version 0.7.15 [16] using standard parameters, except that the maximum insert length (`−a`) was set to 2 Mbp to allow pairs to map on different exons, and that insert size detection was disabled (`−A`). Extra header records (for SQ: AS and for RG: CN, ID, LB, PU, SM, and PL) were added to ease processing and tracking. The resulting BWA SAM formatted alignments were then converted to BAM format, and unmapped as well as non-properly paired CAGE reads were discarded (flag `0×42`). The resulting “CAGEscan pairs” provide individual experimental information on the association of a single-nucleotide-resolution TSS with the body of a gene product.

The CAGEscan pairs were then converted to BED12 format using the program `pairedBamToBed12` version 1.2, in which the score field is the sum of the mapping qualities of each read of the pair. They were then assembled into CAGEscan clusters using the `CAGEscan-Clustering` script version 1.2 and the Phase 1 + 2 FANTOM5 DPI CAGE peaks as seeds. The CAGEscan-Clustering script also takes advantage of the BED12 format, reporting the number of CAGEscan paired-end reads used to assemble each cluster via the score field and the name and position of the seeding CAGE peak via the name, thickStart and thickEnd fields respectively. Finally, the CAGEscan clusters from all libraries were then combined into a single global assembly of “meta-clusters” using the same program and output in BED12 files where the score indicates the number of libraries contributing data to each meta-cluster.

### Code availability

The MOIRAI workflow template used to process the libraries is available as a supplemental XML file on Figshare (DOI: 10.6084/m9.figshare.4792666). MOIRAI enabled the design of a complete data processing pipeline based on the following softwares: FASTX-Toolkit (http://hannonlab.cshl.edu/fastx_toolkit/), TagDust 1.13 [15], rRNAdust 1.03 (http://fantom.gsc.riken.jp/5/sstar/Protocols:rRNAdust) (note that for new projects, we recommend TagDust 2 instead of TagDust 1 and rRNAdust), BWA 0.7.15-r1140 ([16]), SAMtools 0.1.19-44428cd ([17]), pairedBAMtoBED12 1.2 (https://github.com/Population-Transcriptomics/pairedBamToBed12, DOI: 10.6084/m9.figshare.4792672), CAGEscan-Clustering.pl 1.2 (https://github.com/nicolas-bertin/CAGEscan-Clustering, DOI: 10.6084/m9.figshare.4792675) and promexinstats.sh for the annotation (see supplemental material). The software above and standard Unix tools are sufficient to re-implement the pipeline in a different workflow system.

## Data Records

Each CAGEscan library is described with a Sample and Data Relationship Format (SDRF) record, together with the rest of the FANTOM5 data ([9]). For each library, raw sequences in FASTQ format, alignment data in BAM format (including unmapped reads), CAGEscan pairs in BED12 format, CAGEscan clusters in BED12 format and alignment statistics in plain text tabulation-delimited triples (subject, predicate, object), are available in the FANTOM5 data repository. The raw sequences have also been deposited to DDBJ DRA (Data Citation 2).

## Technical Validation

We derived individual library alignment statistics from the MOIRAI data processing pipeline (see Table 1 and Figures 2 and 3A). The statistics count the number of reads discarded at key steps of the processing. “Unextracted” are pairs where the linker was not found, “Artefacts” are pairs that matched the artifact library, or had a low complexity, “rDNA” are pairs that matched the reference rDNA locus (including rRNAs and their spacer regions), “Non-aligned” are pairs where one or both mates were not aligned to the genome, and “Non-proper” are pairs where the mates were not aligned in head-to-head orientation within 2 Mbp. “Duplicates” are the pairs removed during the deduplication step. That is, when there are n pairs with identical coordinates, 1 is kept and *n* − 1 are discarded as “Duplicates”. These statistics show that the amount of PCR duplicates was not larger than the number of CAGEscan pairs, suggesting that the libraries prepared in this study have not been fully exhausted by sequencing.

The library alignment statistics, as well as statistics describing the distribution of CAGEscan TSSs on GENCODE 19 annotations (Figure 3B), also suggest that the biological nature of the samples (cancer cell lines, primary cells, tissue samples and brain tissue) strongly influenced the performance of the CAGEscan protocol used in this study. Albeit displaying the best performance in terms of alignment (largest fraction of CAGEscan pairs), brain tissue derived samples had the lowest rate of known promoters overlapping start sites, hinting at a much greater diversity of alternative promoters usage in human brain. However, since, in this study, all brain tissue derived samples were taken from a single donor, this observation may result from technical batch effect rather than being a general feature of the nature of human brain transcriptome.

## Usage Notes

We have seeded the CAGEscan clustering with FANTOM5 CAGE-defined core promoter regions, however alternative seeding strategies could be envisioned. The 5′ ends of the CAGEscan pairs themselves could be clustered by peak calling and used as a seed, which is the default mode of operation of the `pairedBamToBed12` tool. Foregoing the discovery of alternative promoters, CAGEscan clusters could also be seeded using promoter regions defined by GENCODE models. To discover potential enhancer-associated non-coding RNAs, region corresponding to FANTOM5 enhancers [18] could also be used.

We used a simple alignment strategy that did not take splicing into account. Thus, pairs overlapping splice junctions could not be mapped and CAGEscan clusters lack coverage at the beginning and end of each exon, but this only mildly impacts the main purpose of the method. In addition, since the CAGEscan pairs are anchored at the 5′ end of the transcripts, splice junctions occurring close to the TSS may render some whole loci unmappable. Indeed, transcripts databases such as GENCODE reveal splice junctions very near to the TSS. Trimming the CAGE reads to 20 nt rescued some loci, but other loci were lost due to the decrease of alignment stringency (data not shown). Thus, the development of a spliced alignment workflow would increase the accuracy of our method.

**Figure 3:**
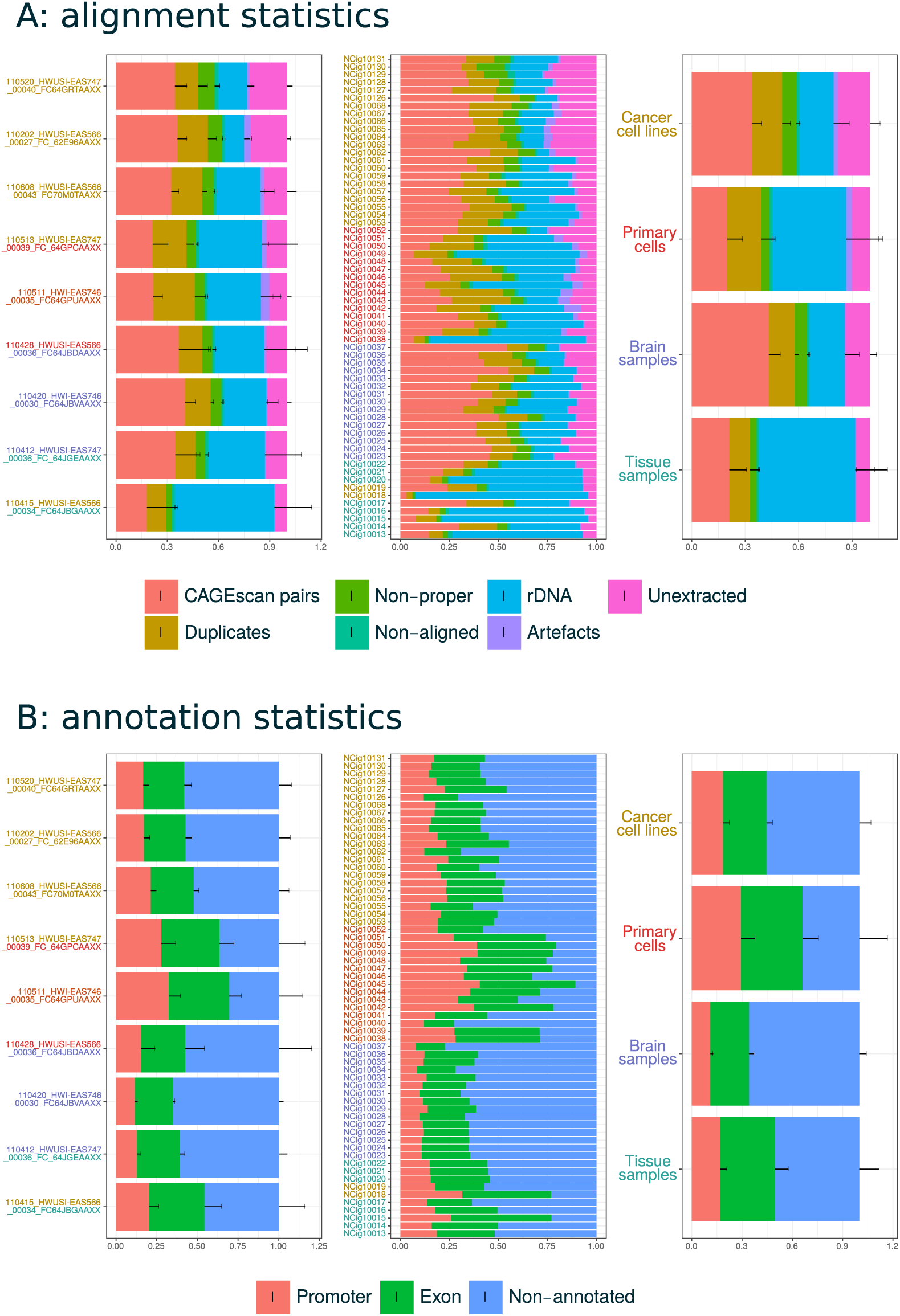
Quality control statistics. A: Fraction of pairs passing all filters (CAGEscan pairs) or discarded at key steps of the processing pipeline (see Figure 2). The central block of stack bars represents each library individually. The left block aggregates them by sequencing batch, named by the sequencing run identifier. The right block aggregates the libraries by sample type. Each sample type is represented by one color, that is also used to color the library identifiers and the sequence identifiers in the other blocks. Batches comprising multiple types are indicated by multiple colors. B: Fraction of pairs starting in a Promoter, Exon, or Other (non-promoter, non-exon) region.

One of the most striking differences between the HeliScopeCAGE-based FANTOM5 CAGE data and the nanoCAGE-based FANTOM5 CAGEscan data is a larger amount of start sites in the gene body, far from the promoter. This can be explained by the lower stringency of the nanoCAGE protocol, which uses template-switching for capturing 5′ ends from limiting amounts of samples [6], where the HeliScopeCAGE protocol, that uses CAP Trapper [19], would not be possible. Readers curious about the position of the random priming site, indicated by the end position of the CAGEscan pairs, will notice that their distribution is very far from random. Control experiments performed using different batches of random primers ordered by different makers confirmed that the quality of the oligonucleotides was not in question (data not shown). In the latest version of the nanoCAGE protocol [20], this problem was solved by the fragmentation of the cDNAs by the “tagmentation” method. Altogether, we recommend to use our latest protocol for making new libraries.

In this study, the CAGEscan libraries were prepared using the nanoCAGE method, but the CAGEscan workflow, which can use any paired-end sequencing of CAGE libraries were the 3′ sequencing read is at a random position in the cDNA, can be applied to other publicly available dataset, for instance made with the RAMPAGE method [21].

## Acknowledgments

FANTOM5 was made possible by research grants for the RIKEN Omics Science Center and the Innovative Cell Biology by Innovative Technology (Cell Innovation Program) from the MEXT to Y.H. It was also supported by research grants for the RIKEN Preventive Medicine and Diagnosis Innovation Program (RIKEN PMI) to Y.H. and the RIKEN Centre for Life Science Technologies, Division of Genomic Technologies (RIKEN CLST (DGT)) from the MEXT, Japan. A.R.R.F. is supported by a Senior Cancer Research Fellowship from the Cancer Research Trust, the MACA Ride to Conquer Cancer and the Australian Research Council’s Discovery Projects funding scheme (DP160101960). We thank RIKEN GeNAS for generation of the CAGEscan libraries, the Netherlands Brain Bank for brain materials, and the RIKEN BioResource Centre for providing cell lines.

## Data Citations

1. FANTOM5 CAGEscan view on the ZENBU genome browser: http://fantom.gsc.riken.jp/zenbu/gLyphs/#config=ZkJi4RdBAFhnsudxePrZxD
2. DDBJ Sequence Read Archive, DRA005606 (2017).

